# Generating whole-brain neural activity and behavior through unified latent dynamics

**DOI:** 10.64898/2026.06.05.730482

**Authors:** Davide Nuzzi, Maurizio Mattia, Giovanni Pezzulo

**Affiliations:** Institute of Cognitive Sciences and Technologies, National Research Council, Rome, Italy; Department of Neuroscience, Istituto Superiore di Sanità, Rome, Italy

**Keywords:** whole-brain activity, generative model, digital twin, behavior simulation, *C. elegans*

## Abstract

Understanding how high-dimensional neural activity and behavior emerge from shared underlying dynamics remains a fundamental challenge in neuroscience. Addressing this problem is key to enabling digital twins that can faithfully reproduce and predict the multiscale brain–behavior dynamics of living systems. Here we present NEBULA (NEural and Behavioral modeling through Unified LAtent dynamics), a generative framework that jointly models whole-brain neural activity and behavior. By applying NEBULA to brain-wide recordings from *C. elegans*, we identify a shared low-dimensional dynamical manifold that underlies both neural activity and behavior, enabling long-horizon generation and targeted in silico interventions. Perturbations of the learned dynamics reveal behaviorally relevant transition points, whereas steering interventions enable controlled manipulation of neural and behavioral states without retraining. These results establish a framework for linking brain dynamics to behavior in a living organism and provide a foundation for scalable virtual experimentation in neuroscience.

## 1 Introduction

In neuroscience, the increasing availability of large-scale neural and behavioral datasets has highlighted the importance of computational models — or brain digital twins — capable of integrating these data to enable large-scale simulations and virtual interventions that emulate realistic experimental manipulations in animals (Wang et al., 2024; Mathis and Mathis, 2026; Angelaki et al., 2026; Patow et al., 2024).

However, developing a computational model that jointly captures brain activity and naturalistic behavior, while also enabling long-horizon generative prediction and virtual interventions, remains a major challenge. State-of-the-art methods that derive low-dimensional manifolds from neural activity typically focus on representation learning, rather than on developing models with robust generative capabilities (Schneider et al., 2023; Kumar et al., 2023). Similarly, state-of-the-art models of human brain activity — whether “foundation models” based on deep learning approaches (D’Angelo and Jirsa, 2022; Millet et al., 2022; Hashemi et al., 2025; Wang et al., 2025b; Jiang et al., 2024; Wang et al., 2025a; Ortega Caro et al., 2024; Schneider et al., 2023; d’Ascoli et al., 2026) or biophysically inspired digital twins (Monteverdi et al., 2023; Hashemi et al., 2025; Alboré et al., 2025; Di Antonio et al., 2026; Kringelbach et al., 2024; Jirsa et al., 2023; Morrison and Young, 2025) — primarily aim to predict neural activity and the effects of virtual lesions, rather than to link neural dynamics and behavior. Other studies have simulated neural population dynamics in *C. elegans* using a variety of approaches — including dynamical systems, switching linear systems, and maximum entropy models — sometimes incorporating biophysical constraints such as the connectome (Kunert et al., 2014; Kunert-Graf et al., 2017; Kunert et al., 2017; Chen et al., 2019; Morrison et al., 2021; Linderman et al., 2019). While these approaches capture important aspects of neurodynamics, they generally treat neural activity in isolation from physical kinematics. Related efforts have also explored whole-body physics-based simulations of animal behavior, including biomechanical models of the fruit fly (Vaxenburg et al., 2025) and virtual rodents controlled by neural networks (Aldarondo et al., 2024). Nevertheless, establishing mechanistic links between brain activity and behavior remains a major challenge, particularly in ways that generalize beyond the distributions encountered during training and support virtual interventions capable of generating realistic neural and behavioral dynamics in previously unseen conditions.

To address this challenge, we introduce NEBULA (NEural and Behavioral modeling through Unified LAtent dynamics), a generative framework that discovers a shared, low-dimensional latent dynamical structure underlying whole-brain activity and behavior. By learning this common dynamical substrate directly from neural recordings, the framework unifies neural and behavioral dynamics within a single probabilistic model, enabling long-horizon generation and targeted counterfactual interventions. We adopt the framework to characterize locomotion behavior and associated whole-brain neural activity in the nematode *C. elegans* (Kato et al., 2015).

We evaluate the framework across several tasks. First, in the encoding setting, NEBULA learns a joint latent manifold of neural and behavioral dynamics. Second, through generative rollouts, it produces realistic long-timescale trajectories of both behavior and neural activity. Third, in the intervention setting, the model enables virtual perturbations that steer both latent model dynamics and behavior toward desired dynamical regimes. These results demonstrate the model’s ability to jointly capture, generate, and control coupled brain–behavior dynamics within a unified framework.

## 2 Results

### 2.1 Characterization of empirical data and virtual kinematic reconstruction

To establish a quantitative foundation for our generative modeling framework, we first characterized the empirical neural and behavioral dataset (see Methods 4.1). We analyzed whole-brain calcium imaging recordings of an immobilized *C. elegans* subject originally acquired by Kato et al. (2015), which capture the simultaneous, continuous activity of 109 neurons (Figure 1a). Although the animal was physically restrained, specific neurons exhibited activity patterns that reliably differentiated distinct fictive locomotor states (“forward”, “slow”, “reverse 1”, “reverse 2”, “reverse sus”, “dorsal turn”, and “ventral turn”). For example, the activity of the AVA command interneuron varied systematically across various locomotor behaviors, enabling their accurate discrimination (Figure 1b). Principal component analysis (PCA) of the full multi-neuron timeseries demonstrates that the global neural dynamics occupies a low-dimensional latent space, wherein the latent state trajectory naturally clusters according to seven locomotor behaviors (Figure 1c).

**Figure 1:**
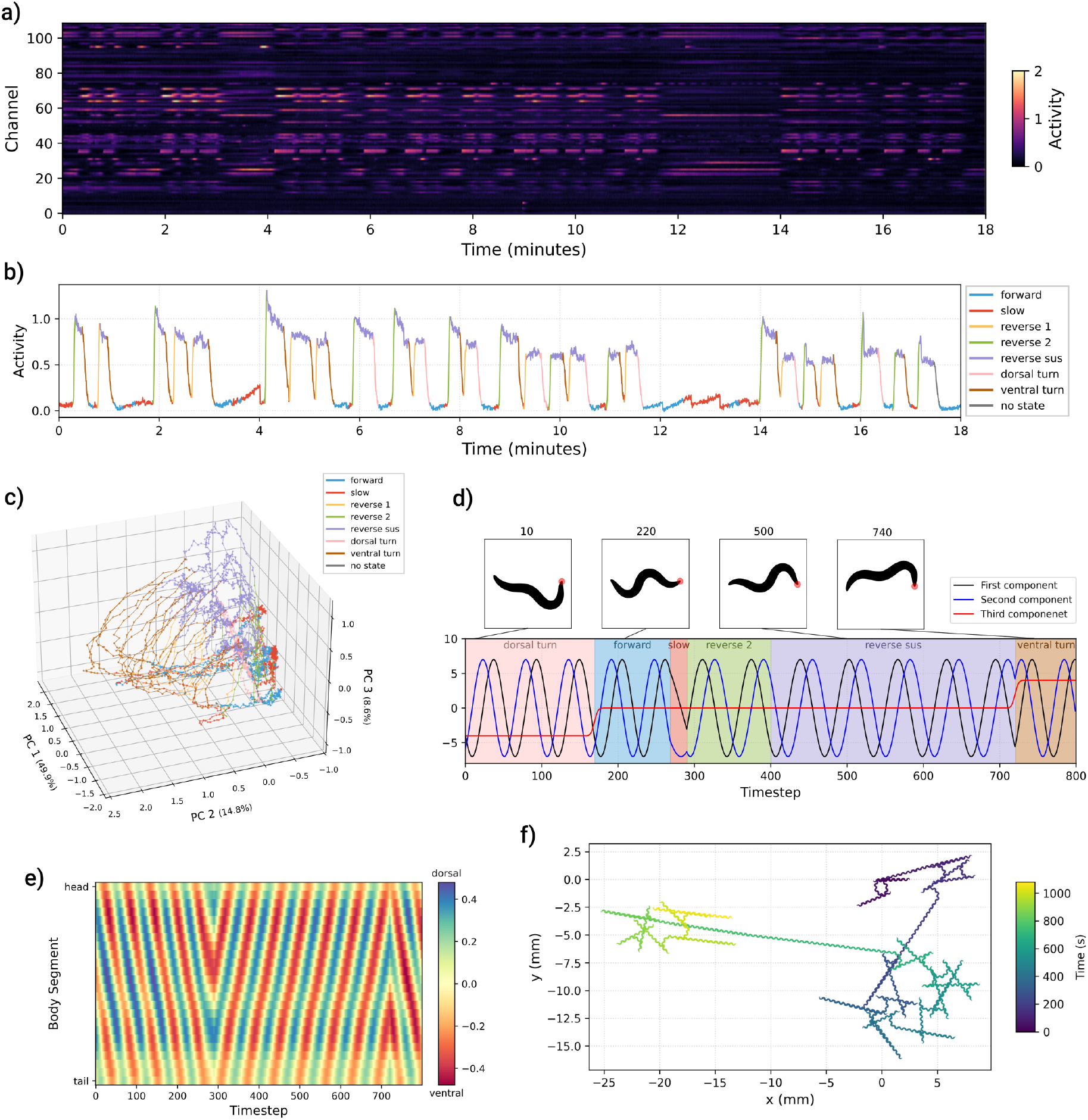
Characterization of empirical data and virtual kinematic reconstruction. **a)** Calcium signals recorded in (Kato et al., 2015) for each single neurons (109 rows). **b)** Trace of the AVA neuron colored according to the behavioral state labelled in (Kato et al., 2015). **c)** PCA of the neural signals. Only the first three components are shown. Each point is colored by the corresponding behavioral state. The percentage of explained variance is shown near each axis label. **d)** Postural reconstruction from discrete behavioral states. The bottom plot shows the temporal evolution of the first three postural components (eigenworm amplitudes): first (black), second (blue), and third (red). Shaded regions denote discrete behavioral states over a 25 seconds window. Top panels display representative reconstructed body shapes at *t* = 10, 220, 500 and 740, with red circles indicating the head position. **e)** Kymogram of body curvature. Postural dynamics reconstructed from the eigenworm amplitudes over the same period as panel **d.** The heatmap illustrates the relative angle (radians) between 25 adjacent body segments from head (top) to tail (bottom). Blue and red colors denote dorsal and ventral bending, respectively. **f)** Simulated spatial trajectory. The path of the worm in the *xy*-plane reconstructed from the sequence of behavioral states over the full 18-minute recording. The trace is colored by elapsed time (seconds), as indicated by the colorbar.

Because the original recording of Kato et al. (2015) inherently lacks physical tracking data, we implemented a kinematic reconstruction pipeline to bridge the discrete fictive locomotor behaviors and continuous spatial movements of the worm (see Methods 4.4). We mapped the sequence of behavioral labels into dynamic body postures represented by linear combinations of the first three principal “eigenworms” (Stephens et al., 2008). By deterministically modulating the phase and amplitude of these postural components based on the active locomotor behavior, we generated a continuous, temporally smoothed sequence of body shapes (Figure 1d). This ego-centric postural evolution is visualized as a kymogram, illustrating the relative angular curvature across 25 adjacent body segments over time (Figure 1e).

Finally, to translate these relative postural changes into ecologically valid spatial locomotion, we processed the synthetic kymograms through the ElegansBot (Chung et al., 2024) biomechanical simulation environment (see Methods 4.4). The simulation resolves the dynamic physical forces between the undulating body segments and a virtual substrate, converting the ego-centric shape changes into absolute 2D spatial coordinates. This pipeline successfully yields a full, continuous spatial trajectory of the virtual worm directly resulting from the fictive behaviors in the original 18-minute empirical recording (Figure 1f), establishing a physical kinematic baseline for evaluating the subsequent generative model outputs.

### 2.2 Neural and behavioral modeling

To capture the complex temporal dependencies linking neural activity and behavior, we developed NEBULA: a probabilistic deep generative framework based on variational state-space modeling (see Methods 4.2). Briefly, the framework consists of three core components: an encoder that compresses the causal history of high-dimensional neural activity into a continuous, low-dimensional latent trajectory; a stochastic transition rate function that models the forward temporal evolution of these latent states; and recurrent observation models (decoders) that project the latent trajectory back into the empirical neural and behavioral data spaces. By jointly optimizing these components, the model learns a temporally predictive and dynamically consistent latent manifold that encapsulates the animal’s motor repertoire.

We applied the framework to the neurobehavioral dataset of Kato et al. (2015), using a three-dimensional continuous latent space. This low-dimensional representation was sufficient to accurately reconstruct both neural and behavioral dynamics while enabling direct visualization of the learned manifold, as each axis corresponds to one of the latent dimensions. Projecting the training dataset into this space via the encoder revealed a highly structured, continuous manifold (Figure 2a), exhibiting a substantially richer repertoire of cyclic trajectories than those observed in behavior alone (Ahamed et al., 2021). When the encoded latent coordinates were colored according to the simultaneous output of the behavioral decoder, the manifold naturally organized into distinct regions associated with different locomotion behaviors.

**Figure 2:**
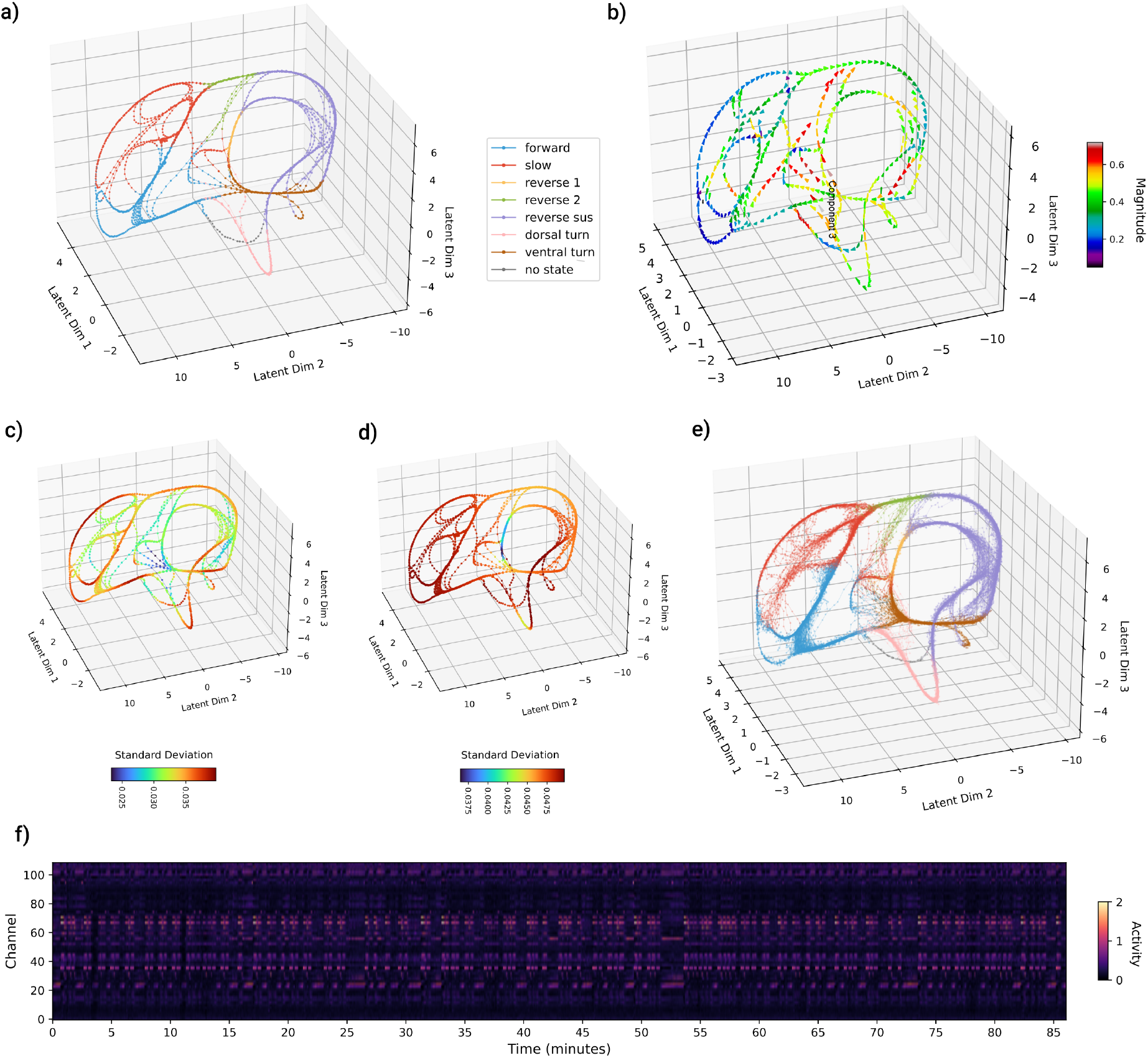
Neural and behavioral modeling through NEBULA. **a)** Encoded latent manifold. The training dataset represented within the three-dimensional latent space of the encoder. Each point is colored according to its ground-truth behavioral label. **b)** Latent transition rates. Arrows indicate the direction and magnitude (color) of the vector field corresponding to the transition rates, visualized only at points on the manifold from panel **a** for clarity. **c)** Encoder standard deviation. Manifold points colored by the standard deviation of the encoder’s output distribution. **d)** Transition standard deviation. Manifold points colored by the standard deviation of the transition rates. **e)** Generative rollout. Latent trajectories produced by the transition module over 30000 timesteps within the three-dimensional manifold. Points are colored by the decoded behavioral state. **f)** Decoded neural activity. Heatmap showing the activity across all neural channels decoded from the generative rollout trajectories over the first 15000 timesteps (≈ 85 minutes). Color indicates simulated activity magnitude.

The temporal evolution along this manifold is governed by the learned transition rates. By evaluating these transition rates across the visited latent states, we can visualize the vector field that drives the system’s dynamics, where arrows indicate the instantaneous magnitude and direction of the mean temporal derivatives (Figure 2b). Furthermore, being the framework doubly probabilistic, it yields localized, state-dependent estimations of uncertainty for both the inference and generation processes. Mapping the standard deviation of the distributions across the visited manifold reveals that the encoder and the transition rates exhibit entirely distinct spatial profiles of variance. The uncertainty of the encoder (Figure 2c) is highly localized, reaching its maximum values near the centers of large, stable behavioral arcs, away from critical trajectory bifurcations or sub-loops. Conversely, the intrinsic stochasticity of the transition rates (Figure 2d) exhibits less overall variation across the manifold but is noticeably elevated within the specific lobe corresponding to forward and slow locomotion, far from the origin of the latent space.

To test the stability and autonomy of the local dynamics learned by the model, we decoupled it from the empirical data and generated a long-horizon, unsupervised rollout of 30000 timesteps, corresponding to roughly 168 minutes (see Methods 4.2). Guided solely by autoregressive sampling from the transition rates, the generated trajectory remains dynamically stable. The generative rollout accurately reproduces the topological structure, cyclical patterns, and behavioral partitioning of the original training data without collapsing or diverging (Figure 2e), thus characterizing this manifold as an attractor. For visual clarity regarding the underlying structural geometry, the plotted trajectories in these low-dimensional latent space represent the sequence of distribution means (*µ*) rather than the raw stochastic samples (*z*). Simultaneously, the framework decodes continuous neural activity from this same autonomous latent trajectory. Over the roughly 168-minute simulated duration, the generated neural timeseries exhibits complex, non-periodic population dynamics that remain stable without collapsing to a single equilibrium point (Figure 2f).

### 2.3 Quantitative assessment of the model’s generative dynamics

To quantitatively determine whether the generative rollouts of NEBULA faithfully capture the statistical structure of the biological data, we compared the macroscopic properties of the generated activity in Figure 2e against the empirical training dataset. We first evaluated the accuracy of the behavioral decoding. The framework accurately reproduced the global behavioral state frequencies observed in the original data (Figure 3a), yielding a Kullback-Leibler (KL) divergence of 2.7 · 10^−3^. To statistically evaluate whether the generated rollouts originate from the same underlying distribution as the biological data, we conducted a Pearson’s *χ*^2^ test of homogeneity across both independent samples. The generative dynamics were found to be statistically indistinguishable from the empirical data (*χ*^2^ = 11.06, *p* = 0.136). Beyond static frequencies, the generative rollout successfully internalized the temporal syntax of the animal’s motor repertoire, demonstrating a high degree of concordance between the empirical and generated behavioral transition matrices (Figure 3b; KL divergence = 6.3· 10^−3^). To confirm this alignment, we tested the homogeneity of the Markov chains. Independent Pearson’s *χ*^2^ tests revealed no significant deviations for any individual state’s outbound transitions (*p >* 0.05 for all rows). Furthermore, an aggregated global test confirmed that the overarching empirical and generated transition matrices remain statistically indistinguishable (*χ*^2^ = 5.96, *p* = 0.999).

**Figure 3:**
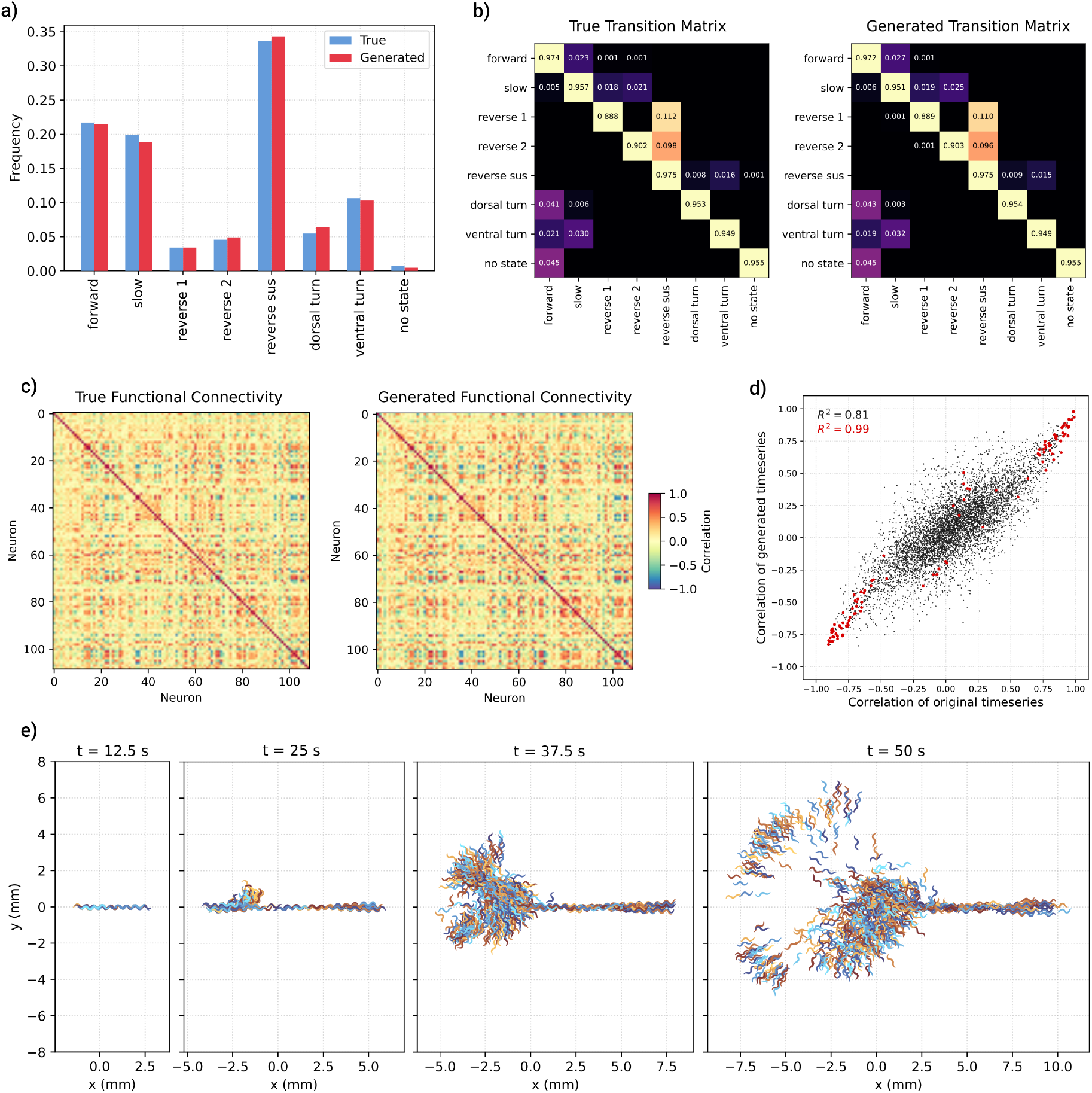
Quantitative validation of generated neural and behavioral dynamics. **a)** Behavioral state frequencies. Comparison between the true distribution of behaviors from the training set and the distribution decoded from the 30000 timesteps generative rollout visualized in Figure 2e. **b)** Behavioral transition matrices. Comparison of transition probabilities between states in the training data (left) and the decoded generative rollout (right). Values denote non-zero probabilities. **c)** Functional connectivity matrices. Pairwise linear correlations between all neural channels for the original data (left) and those decoded from the generative rollout (right). **d)** Correlation between real and generated functional connectivity. Comparison of functional connectivity (computed via pairwise linear correlations) between the original neural timeseries and that decoded from generated latent trajectories. Each point represents a unique pair of neural channels. Black points include all channel pairs, while red points denote a subset of consistent neurons identifiable across all worms (see text and (Kato et al., 2015)). The data for the comparisons are illustrated in the functional connectivity matrices of panel **c. e)** Stochastic divergent trajectories. Snapshots at four time points (*t* = 12.5, 25, 37.5, 50 s) showing the spatial dispersal of 500 simulated worms. All simulations begin from the same latent state and spatial origin. The diversity in trajectory and body posture is driven by the stochasticity of the latent transition rates.

Because the continuous neural activity decoded from the generative rollout (Figure 2f) is a novel stochastic realization governed by the latent transition prior, the resulting timeseries naturally diverges from the exact temporal sequence of the original 15-minute empirical recording. Therefore, rather than comparing direct temporal alignments, we quantified the fidelity of the generated neural dynamics by assessing the preservation of statistical features like the functional connectivity. We computed the pairwise Pearson correlation matrices for all 109 neurons across both the true and generated datasets (Figure 3c). Drawing on the analytical framework of Representational Similarity Analysis (RSA) (Kriegeskorte et al., 2008), we extracted the unique pairwise correlations (upper triangular elements of the functional connectivity matrices) from both empirical and generated time series (Figure 3c), and quantified their similarity by computing the element-wise correlation between these matrices (Figure 3d). The global network topology is highly conserved, demonstrating a strong linear relationship across all neuron pairs (*R*^2^ = 0.81, black points). Interestingly, this functional preservation is most precise within the subset of neurons most involved in locomotion. We isolated a specific subset of 15 extensively characterized command and premotor neurons (including the AVA, AVB, and AIB classes; see Methods 4.1) that were unambiguously identified in the physical recording (Kato et al., 2015). When restricting the comparative analysis to correlation pairs containing at least one of these neurons, the correlation between the empirical and generated network states increases (*R*^2^ = 0.99, red points).

Generated neural trajectories are then statistically equivalent replicas of the experimental ones, even though no uncertainty associated with predicting future states from past neural activity has been incorporated into the generative process. Thus, the widening and compression of trajectory bundles in the generative phase (Figure 2e) can be attributed solely to the state-dependent Markovian dynamics in the latent space determined by the learned transition rates. In other words, the stochastic latent dynamics is self-consistent, and its dimensionality (*D* = 3) likely represents the irreducible manifold explored by C. elegans neural activity. This is also consistent with the dynamics being organized as a finite-state automaton (Berman et al., 2016), a graph of behavioral transition points connected by arcs, since any such graph embeds without self-intersection in three dimensions (Cohen et al., 1997), making *D* = 3 sufficient to represent it faithfully.

Finally, to visualize how the latent transition dynamics translate into physical movement, we simulated the spatial dispersal of an ensemble of virtual worms. We initialized 1000 independent generative rollouts from the same starting condition (*z*_0_, the encoded latent state of the first empirical sample) and tracked their continuous spatial trajectories over a 60-second physical simulation horizon. Snapshots of this ensemble at progressively longer timescales reveal the rich unfolding of the model’s behavioral repertoire (Figure 3e). During the early phases (*t* = 12.5 and 25 s), the trajectories diverge relatively slowly, clustering along the primary longitudinal axis as individual rollouts sample sustained, stable locomotor states (forward or backward propulsion). However, at longer horizons (*t* = 37.5 and 50 s), the accumulated stochasticity within the latent manifold triggers probabilistic bifurcations into turning states. This causes the paths to branch outward, demonstrating the model’s capacity to spontaneously generate diverse, multi-step behavioral sequences. Notably, the final spatial distribution of the ensemble at 50 seconds is highly anisotropic, forming distinct, asymmetric spatial lobes. This physical asymmetry is a macroscopic manifestation of the biological biases encoded within the latent manifold—capturing the unequal transition probabilities between dorsal and ventral turns that were present in the empirical training data.

### 2.4 Robustness of generative dynamics to latent perturbations

To evaluate the structural stability of the learned manifold, we investigated how the generative dynamics respond to varying degrees of stochastic perturbation. We first analyzed the effect of applying continuous, global noise across the entire three-dimensional latent space during 30000-step generative rollouts. The injected noise was sampled from an isotropic multivariate Gaussian distribution with a diagonal covariance matrix, parameterized by a scalar noise intensity (standard deviation).

We quantified the degradation of the model’s output as a function of this noise intensity. For the behavioral output (Figure 4a), we measured the KL divergence between the generated and empirical behavioral state distributions. Notably, the behavioral frequencies remain essentially unchanged until the noise intensity exceeds the endogenous stochasticity set by the autonomous Markovian dynamics by more than a factor of 20. For the neural output (Figure 4b), we computed the Mean Squared Error (MSE) between the functional connectivity matrices of the generated and original neural timeseries. The neural dynamics demonstrate a slightly higher sensitivity, with the MSE lifting above zero around an intensity of 0.3, followed by a steady increase at higher noise levels. Together, these results show that the generative model is highly robust, implementing a stable attractor manifold that preserves its topology and dynamical stability even under continuous perturbations an order of magnitude larger than the stochasticity experienced during training or autonomous generation.

**Figure 4:**
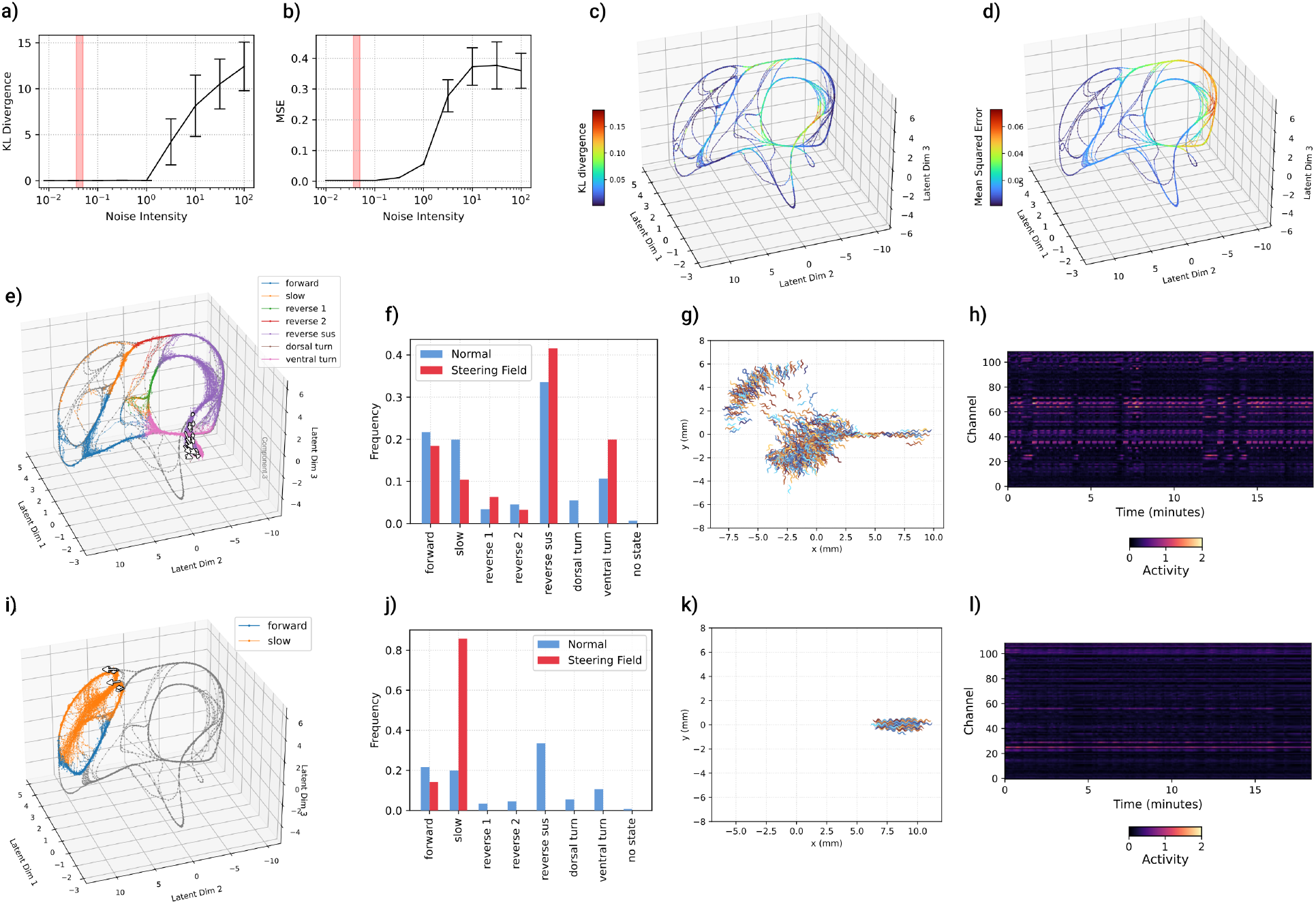
Robustness to latent perturbations and targeted steering of behavioral dynamics. **a)** Behavioral distribution robustness. KL divergence between the behavioral state frequencies of a baseline (noise-free) generative rollout and those subjected to varying intensities of white Gaussian noise. The red shaded region indicates the standard deviation range associated with the learned transition rates. Error bars represent the standard deviation across 20 independent generative samples per intensity level. **b)** Functional connectivity robustness. MSE between the functional connectivity matrices of the baseline generative rollout and those generated under varying intensities of latent noise. The red shaded region indicates the standard deviation range natively expressed by the learned transition rates. Error bars represent the standard deviation across 20 independent generative samples per intensity level. **c)** Behavioral sensitivity to localized noise. KL divergence between the behavioral distributions of a baseline generative rollout and trajectories subjected to localized latent perturbations (*σ* = 5 within radius 1). Color indicates the discrepancy resulting from noise applied at each manifold location. **d)** Connectivity sensitivity to localized noise. MSE between functional connectivity matrices of baseline and locally perturbed rollouts. Color maps represent the local impact of noise on the accuracy of decoded neural interactions. **e-h)** Steering toward ventral turns. Targeted intervention on the transition rates to modulate behavioral priors, specifically to increase the frequency of ventral turns. In panel **e**, baseline manifold is shown in grey, while the steered trajectory is colored by its decoded behavioral state. White arrows represent the learned steering field vectors (magnitude *>* 0.05). Panel **f** shows the behavioral state frequencies, providing a comparison between the distribution of behaviors from the baseline generative rollout and the distribution decoded from the steered generative rollout. Panel **g** shows the spatial dispersal of simulated worms driven by the steered latent transition rates. All simulations begin from the same latent state and spatial origin. Panel **h** shows the activity across all neural channels decoded from the steered generative rollout trajectories over 18 minutes. Color indicates simulated activity magnitude. **i-l)** Steering toward forward and slow locomotion. A second targeted intervention on the transition rates designed to incentivize forward and slow behaviors while suppressing reversals and turns. Panels **i** through **l** mirror the data modalities of panels **e** through **h**, respectively illustrating the steered latent manifold, the corresponding shift in behavioral state frequencies, the resulting spatial dispersal of simulated worms, and the decoded neural activity under this alternative steering policy.

We next examined the sensitivity of different regions of the manifold by applying localized, non-directional noise. We considered 3137 equally spaced points across the encoded empirical manifold and defined a local boundary (a sphere of radius 1) around each coordinate. We then initiated 30000-step generative rollouts, injecting a severe isotropic Gaussian noise (*σ* = 5) strictly when the latent trajectory *z*_*t*_ entered the specific spherical region. The resulting discrepancies were then mapped back onto the originating manifold coordinates as a color gradient.

The spatial sensitivity of the manifold varies significantly depending on the decoded modality. The behavioral sensitivity map (KL divergence, Figure 4c) reveals that the most critical points of failure occur just before major trajectory bifurcations. For instance, high sensitivity is localized in the *reverse sus* region immediately preceding the split into dorsal or ventral turns. A similarly sensitive zone appears along the *reverse 2 arc*, where localized noise is sufficient to kick the latent trajectory against the directed flow of the transition rates, forcing it prematurely back into the forward/slow cycle.

Conversely, the sensitivity map for the neural functional connectivity (MSE, Figure 4d) exhibits a different spatial profile. While the regions corresponding to reversals and turns remain highly susceptible to noise-induced disruption, the region representing forward and slow locomotion exhibits high dynamic robustness, absorbing the high-intensity localized noise with almost negligible impact on the fidelity of the decoded neural signals.

### 2.5 Targeted modulation of behavior via latent manifold steering

While our generative framework simulates spontaneous behavior, investigating goal-directed control requires a mechanism to actively influence these dynamics. To achieve this, we introduced a latent steering module that applies sparse, targeted perturbations to the state-dependent transition rates, pushing the system toward specific behavioral patterns without retraining the underlying manifold (see Methods 4.5).

We first tested the system’s ability to promote a specific turning behavior by explicitly favoring the *ventral turn* state while suppressing all others. A 10000-step generative rollout under this steering policy demonstrates that the model successfully achieves the behavioral goal while maintaining the structural integrity of the continuous latent space (Figure 4e). The steered trajectory (colored) remains closely anchored to the baseline encoded manifold (grey), but the *dorsal turn* arc is completely circumvented. Visualizing the learned steering field (white arrows, filtered for magnitude *>* 0.1) reveals that interventions are highly localized, acting only at critical junction points rather than uniformly across the space. The resulting behavioral frequencies (Figure 4f) confirm the targeted increase in ventral turns and the suppression of dorsal turns. Crucially, this localized intervention triggers a cascading effect across the entire behavioral distribution: because the neural manifold enforces strict transition dynamics, reaching a turn requires transitioning through specific reversal states. Consequently, promoting the ventral turn inherently increases the frequency of the *reverse sus* state and proportionally decreases time spent in *forward* and *slow* locomotion, highlighting the deeply intertwined, complex nature of the neurobehavioral manifold. This targeted behavioral shift physically manifests in the spatial dispersal of the simulated worms (Figure 4g), which, unlike the unsteered baseline, lacks a concentration of trajectories in the bottom-left region (compare with Figure 3e-right). This behavioral modulation is accompanied by the corresponding neural activity decoded directly from the steered latent trajectory (Figure 4h).

In a second experiment, we inverted the objective, promoting only the *forward* and *slow* locomotor states while suppressing all reversals and turns. The resulting rollout (Figure 4i) shows the trajectory effectively trapped within the region where *forward* and *slow* locomotor states are expressed. The learned steering vectors emerge at the bifurcation boundaries leading toward the *reverse 2* state, pointing in direct opposition to the natural dynamical flow to prevent the system from exiting the *forward* – *slow* locomotor cycle. Furthermore, this structural confinement leaves the lowermost arcs of the *forward* region entirely unvisited, indicating that these specific paths represent return trajectories exclusively accessed following turning events. The behavioral distribution (Figure 4j) confirms the near-total suppression of non-locomotor states. Interestingly, while both *forward* and *slow* behaviors were equally incentivized, the dynamical constraint causes *slow* locomotion to become the overwhelmingly dominant state (comprising *>* 80% of the rollout), while the frequency of forward locomotion actually decreases relative to the baseline spontaneous dynamics. Accordingly, the spatial dispersal of the simulated ensemble exhibits minimal lateral spread, resulting in extended, primarily forward trajectories devoid of turning events (Figure 4k), supported by the steady-state neural activity decoded under this constrained steering policy (Figure 4l).

## 3 Discussion

In this study, we aimed to bridge a critical gap in computational neuroscience: the development of a generative digital twin capable of jointly modeling whole-brain neural dynamics and continuous naturalistic behavior (Wang et al., 2024; Mathis and Mathis, 2026; Angelaki et al., 2026; Patow et al., 2024).

To address this challenge, we developed NEBULA (NEural and Behavioral modeling through Unified LAtent dynamics), a generative framework that uncovers a shared, low-dimensional latent dynamical linking neural population activity and behavior. Applying NEBULA to whole-brain recordings from *C. elegans* revealed that diverse locomotor behaviors arise from coherent trajectories within this common, low-dimensional dynamical manifold, which can autonomously generate realistic neural and behavioral dynamics while supporting counterfactual *in-silico* perturbations.

NEBULA maps high-dimensional neural and behavioral time series from *C. elegans* onto a highly structured, continuous low-dimensional manifold (here, three-dimensional) and internalizes the underlying stochastic neurobehavioral dynamics through learned state-dependent transition rates that governs latent-state evolution. As demonstrated in our results, when decoupled from the empirical data, the model can autonomously generate long-horizon synthetic rollouts that remain dynamically stable. These generated trajectories strictly preserve the temporal syntax of the animal’s natural motor repertoire and maintain near-perfect functional connectivity (*R*^2^ = 0.99) among the core command and premotor neurons governing locomotion. Finally, we demonstrated the physical validity of these latent dynamics by translating the discrete behavioral outputs into continuous spatial locomotion using a set of eigenworms as low-dimensional basis (Stephens et al., 2008) and the ElegansBot biomechanical simulator (Chung et al., 2024). This complete generative pipeline, from stochastic latent transitions to simulated 2D spatial trajectories, confirms that the framework successfully captures the mechanistic link between abstract neural population states and physical motion.

Beyond generation, a true digital twin must support counterfactual simulations and targeted interventions. We validated this capability by subjecting the learned manifold to both unstructured perturbations and goal-directed steering. Noise-injection experiments revealed that the generative dynamics are robust to global stochasticity while exhibiting localized sensitivity at trajectory bifurcations, mirroring the biological topology where specific neurobehavioral transitions (such as the initiation of a turn) represent critical dynamical junctions. Furthermore, by introducing a deterministic latent steering module, we demonstrated the ability to enact targeted *in-silico* interventions without retraining the base model. By applying sparse, minimal perturbations to the transition dynamics, we directed the virtual animal to adopt specific behavioral regimes, such as prioritizing ventral turns or remaining confined to forward locomotion. This steering mechanism provides a computational analog to experimental virtual lesions or optogenetic stimulations (Deisseroth, 2015) and extends recent whole-brain modeling efforts that utilize latent space perturbations to discover transitions between dynamic brain states (Sanz Perl et al., 2023; Langdon and Engel, 2025). Together, these results demonstrate that the framework provides a controllable, dynamic environment to test hypotheses about how targeted latent modulations cascade into physical behavior.

Our framework combines the strengths of existing approaches that either focus on representation learning without robust generative capabilities (Schneider et al., 2023; Kumar et al., 2023) or model neural and behavioral dynamics in isolation in *C. elegans* (Kunert et al., 2014; Kunert-Graf et al., 2017; Kunert et al., 2017; Chen et al., 2019; Morrison et al., 2021; Linderman et al., 2019) and other animals (d’Ascoli et al., 2026; D’Angelo and Jirsa, 2022; Hashemi et al., 2025; Wang et al., 2025b; Jiang et al., 2024; Wang et al., 2025a; Ortega Caro et al., 2024; Schneider et al., 2023; Monteverdi et al., 2023; Hashemi et al., 2025; Alboré et al., 2025; Di Antonio et al., 2026; Kringelbach et al., 2024; Jirsa et al., 2023; Morrison and Young, 2025; Aldarondo et al., 2024; Vaxenburg et al., 2025; Durstewitz, 2017; Koppe et al., 2019). The key innovation of NEBULA is the integration of manifold learning and latent transition dynamics within a unified generative framework, enabling robust generation of coupled neural and behavioral trajectories, realistic simulations of behavior, and targeted virtual interventions that steer neural and behavioral states in a controlled manner without retraining. Together, these capabilities establish a digital twin that jointly captures brain and behavior.

While our results demonstrate the stability and flexibility of the NEBULA framework, the current implementation relies on a dataset derived from an immobilized animal exhibiting fictive locomotion (Kato et al., 2015). Consequently, the generated trajectories operate in an open-loop regime, lacking the real-time sensory and proprioceptive feedback that naturally constrains and drives biological movement. Future iterations of this digital twin must integrate multi-modal data from freely moving subjects to explicitly model these closed-loop sensorimotor interactions (Nguyen et al., 2016). Expanding the framework to capture continuous environmental feedback will be essential for modeling the continuous dynamics of planning and acting in naturalistic tasks, and exploring how unsupervised pretraining primes networks for task execution (Zhong et al., 2025). Furthermore, although we demonstrate our framework using neurobehavioral trajectories from *C. elegans*, the framework is general and could be extended in future studies to model the coupled brain–behavior dynamics of other organisms. It can also be used to test theories that describe brain function in terms of generative modeling and statistical inference (Parr et al., 2022; Keller and Mrsic-Flogel, 2018; Knill and Pouget, 2004; Doya, 2007; Pezzulo et al., 2018; Maass et al., 2002; Buonomano and Maass, 2009), by learning generative models directly from data to interpret neural activity and perform virtual experiments. Ultimately, scaling this unified latent space approach to incorporate active sensory loops and behavior in more complex organisms will provide a foundation for advancing theoretical understanding as well as enabling large-scale virtual interventions in neuroscience.

## 4 Methods

### 4.1 Dataset

The empirical data used to train our generative model were obtained from previously published whole-brain calcium imaging experiments of *C. elegans* (Kato et al., 2015), publicly accessible via the Open Science Framework (https://osf.io/2395t/). For our modeling, we utilized a single recording session of a wild-type subject (dataset *TS20140715e_lite-1_punc-31_NLS3_2eggs_56um_1mMTet_basal_1080s* contained in the *WT_NoStim*.*mat* file). During data acquisition, the animal was physically immobilized within a microfluidic chamber to enable stable, single-cell resolution pan-neuronal imaging.

The continuous neural observation matrix, ***x***, comprises the simultaneously recorded activities of 109 neurons. The physiological signal is quantified as the normalized variation in calcium fluorescence (Δ*F/F*), serving as a proxy for intracellular neural activation. This recorded ensemble spans the head ganglia and includes the majority of head motor neurons, sensory neurons, interneurons, and anterior ventral cord motor neurons. A core subset of 15 extensively characterized command and premotor neurons (AIBL, AIBR, ALA, AVAL, AVAR, AVBL, AVER, RID, RIML, RIMR, RMED, RMEL, RMER, VB01, and VB02) were unambiguously identified within this population. This inclusion ensures that the multi-dimensional latent manifold inherently captures the fundamental sensorimotor networks driving locomotion.

The corresponding discrete behavioral sequence, ***y***, consists of “fictive” locomotor states. Because physical locomotion was prevented by immobilization, we utilized the behavioral assignments established by Kato et al. (2015). By validating that specific neural activation patterns in freely moving animals strongly predict kinematic outputs, the original authors successfully assigned a fictive motor command to each temporal frame of the immobilized recording. We adopt this validated annotation scheme directly, classifying the sequence into seven mutually exclusive states: “forward”, “slow”, “reverse 1”, “reverse 2”, “reverse sus”, “dorsal turn”, and “ventral turn”. In our framework, this discrete sequence provides the empirical ground truth for optimizing the behavioral decoder and serves as the driving input for subsequent kinematic simulations.

### 4.2 Variational latent dynamics framework

Let ***x*** denote the continuous multivariate neural recordings, ***y*** denote the corresponding discrete behavioral states, and ***z*** represent the underlying low-dimensional latent variables. Bold variables indicate temporal sequences (e.g., ***z***_0:*T*_ denotes the latent trajectory up to timestep *T*, and *z*_*t*_ specifies the state at time *t*). We formulate a probabilistic generative model parameterized by neural network weights *θ*. The joint probability over a trajectory of length *T* factorizes as:

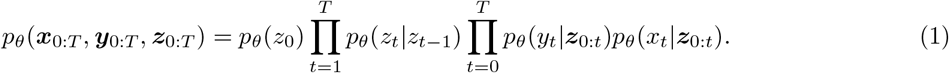

The generative process is initialized with a standard isotropic Gaussian prior, *p*_*θ*_(*z*_0_) = *N* (*z*_0_; 0, I). Subsequent latent evolution follows a stochastic Markovian process defined by the transition rate function *p*_*θ*_(*z*_*t*_|*z*_*t*−1_) = *N* (*z*_*t*_; *µ*_*θ*_(*z*_*t*−1_),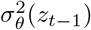). To stabilize optimization and ensure bounded latent trajectories, the transition mean is parameterized as a residual update: *µ*_*θ*_(*z*_*t*−1_) = *z*_*t*−1_ + *α* tanh(*f*_*θ*_(*z*_*t*−1_)). Here, *f*_*θ*_ is a non-linear mapping realized by a neural network, the hyperbolic tangent restricts the maximal per-step deviation, and *α* is a learnable scaling factor determining the transition magnitude. During optimization, latent states *z*_*t*_ are differentiably sampled using the standard reparameterization trick.

The observation models map the latent trajectories to the data spaces. Unlike the Markovian state transitions, the decoding of both modalities at time *t* depends on the entire latent history ***z***_0:*t*_, implemented via recurrent neural networks (RNNs). This recurrent architecture allows the decoders to integrate information across longer temporal scales than the immediate Markovian latent transitions, thereby capturing the long-range dependencies inherent in both neural and behavioral dynamics. The behavioral decoder outputs the logits of a categorical distribution over the discrete states *y*_*t*_. Simultaneously, the neural decoder predicts the continuous signals *x*_*t*_, serving as the mean of a Gaussian emission distribution with unit standard deviation.

Because exact inference in this non-linear generative model is computationally intractable, we employ amortized variational inference, framing the system as a deep variational state-space model. We introduce an amortized variational posterior, *q*_*ϕ*_, which serves as the encoder mapping the high-dimensional neural data into the continuous latent space. The joint posterior over the full latent trajectory is factorized as:

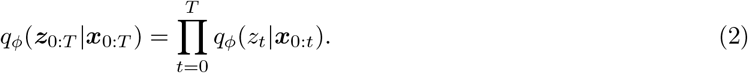

This formulation restricts the latent state estimate *z*_*t*_ strictly to the causal history of the neural signals up to time *t*. By constraining the encoder to act as a filtering distribution that precludes non-causal access to future observations, we enforce the discovery of a temporally predictive and dynamically consistent latent manifold. Architecturally, the encoder is parameterized by a recurrent neural network that outputs the parameters (mean and diagonal covariance) of a Gaussian distribution at each timestep. The latent representations *z*_*t*_ are subsequently drawn using the standard reparameterization trick to maintain end-to-end differentiability during optimization.

To optimize the model parameters, we maximize the log-evidence (marginal likelihood) of the observed sequences. The exact marginal likelihood is obtained by integrating out the latent variables from the joint distribution:

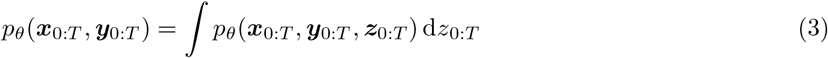

By introducing the variational posterior, we can rewrite the marginal likelihood as an expectation over the latent trajectories. Dropping the time subscripts for notational brevity this becomes:

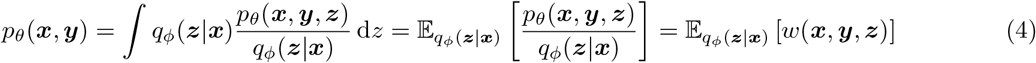

where *w*(***x, y, z***) represents the importance weight of a given trajectory. Standard optimization of the Evidence Lower Bound (ELBO) using a single trajectory sample can suffer from instability and representation collapse during long autoregressive generative rollouts. To obtain a tighter and more robust bound, we adopt an Importance-Weighted Autoencoder (IWAE) framework. We draw *K* independent latent trajectories ***z***^(*k*)^ from the filtering encoder, with the expectation taken over the product distribution 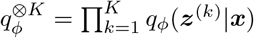. Applying Jensen’s inequality to the logarithm of the *K*-sample expected value yields the importance-weighted lower bound:

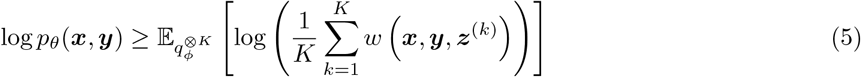

Due to the concavity of the logarithm, this multi-sample bound is strictly closer to the true log-evidence than the standard ELBO. For numerical stability during gradient descent optimization, the objective we maximize is computed using the log-sum-exp formulation:

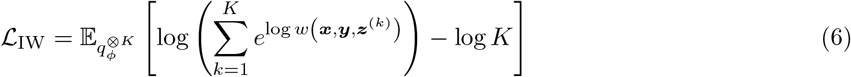

The explicit log-weight of a trajectory decomposes sequentially into the initial prior divergence, the transition dynamics divergence, and the observation likelihoods for both modalities:

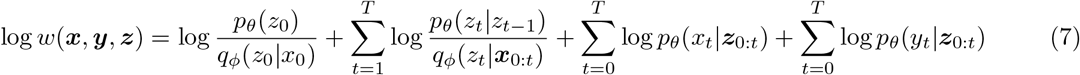

Finally, to ensure the encoded latent states are adequately spaced out across the manifold and to prevent representational crowding, we augment the final optimization objective with a repulsive regularization term applied to the latent states:

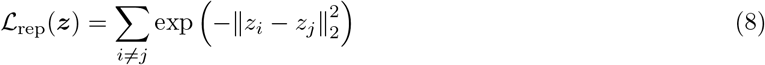

While such structural manifold constraints could theoretically be embedded within a more complex initial prior *p*(*z*_0_), enforcing them via an explicit auxiliary penalty keeps the core variational formulation highly tractable and simplifies the optimization dynamics. During training we introduce scalar weighting coefficients to each constituent term within the log-weight function and to the repulsive loss term to balance the relative influence of the prior, transition dynamics, multimodal observation likelihoods and repulsion force during optimization.

To optimize the model, the algorithmic forward pass strictly separates the encoding of latent states from the evaluation of the transition dynamics. During training, latent trajectories ***z***_0:*T*_ are sampled entirely through the filtering encoder *q*_*ϕ*_ conditioned on the empirical data ***x***_0:*T*_. The transition model *p*_*θ*_ is never used to sample states during optimization. Instead, for each encoded state *z*_*t*−1_, the transition network computes the Gaussian parameters of the conditional prior *p*_*θ*_(*z*_*t*_|*z*_*t*−1_). These parameters are then evaluated against the subsequent encoded state *z*_*t*_ to compute the transition likelihood within the importance weight. This one-step evaluation prevents the accumulation of recursive errors during optimization, effectively acting as a data-driven forcing mechanism that grounds the transition rates directly in the encoded manifold. Once optimized, the model is decoupled from the data and deployed as an autoregressive generative system. Starting from an initial condition *z*_0_, long-horizon latent trajectories are synthesized entirely in the latent space by recursively sampling from the learned transition rate function, *z*_*t*_ ~ *p*_*θ*_(*z*_*t*_|*z*_*t*−1_). Despite the transition rates being trained exclusively on single-step conditional targets, these autoregressive rollouts remain dynamically stable over extended timescales. Finally, these stochastically generated latent trajectories are passed through the recurrent observation models to emit the corresponding synthetic neural signals and behavioral sequences, allowing for the autonomous generation of dynamically consistent neurobehavioral rollouts.

### 4.3 Network architecture and optimization details

Across all neural network modules, the internal hidden dimension was fixed to 128, and the continuous latent space was defined with a dimensionality of 3. The filtering encoder *q*_*ϕ*_ is composed of an initial linear mapping followed by three residual blocks, terminating in a Gaussian head that outputs the mean and diagonal covariance of the latent state. The non-linear mapping *f*_*θ*_ within the transition model *p*_*θ*_ mirrors this architecture exactly, consisting of a linear projection, three residual blocks, and a final layer that parameterizes the Gaussian transition distribution. To capture temporal dependencies, both decoders utilize Gated Recurrent Unit (GRU) networks. The behavioral decoder consists of a single GRU layer that projects directly to the discrete state logits. The neural decoder is more expressive, passing the output of a GRU layer through two additional residual blocks before a final linear readout generates the continuous neural predictions. The model was optimized over 5000 iterations using the Adam optimizer with a learning rate of 3 × 10^−4^. During training, we sampled mini-batches of 256 trajectory segments, each with a fixed temporal window of *T* = 128 timesteps. Note that providing a history of past observations is crucial for enabling the model to learn generative capabilities. In a control experiment with the temporal window set to *T* = 1, the model recovers a sufficiently differentiated internal manifold but does not support generative rollouts (Supplementary Figure S1).

For the estimation of the importance-weighted bound, we drew *K* = 5 independent latent trajectories per sequence. Finally, to balance the constituent terms of the objective function, the following scalar weights were applied: 1 × 10^−5^ for the initial prior divergence, 1 for the transition dynamics divergence, 1 for the neural observation likelihood, 5 for the behavioral observation likelihood, and 5 for the auxiliary repulsive regularization term.

### 4.4 Kinematic reconstruction via eigenworm projection

To translate the discrete sequences of generated behavioral labels into continuous, visually interpretable locomotion, we map the model’s state outputs onto a virtual worm model. This is achieved using the established “eigenworms” framework (Stephens et al., 2008) which demonstrates that the vast majority of C. elegans postural variance can be captured by a low-dimensional basis set of postural modes. Any specific body shape can be expressed as a linear combination of these basis vectors and their time-varying scalar coefficients.

For our kinematic reconstruction, we restricted the basis set to the first three principal eigenworms. Through heuristic evaluation of empirical video recordings, we found that modulating just these three primary components was sufficient to capture the core behavioral repertoire, yielding sequences of body postures that visually align with expected biological kinematics.

To ensure realistic, continuous motion and prevent disjointed transitions when the behavioral label changes, the postural dynamics are governed by a continuous phase variable, *ϕ*_*t*_, which accumulates over time. The first two eigenworm coefficients, *c*_1_ and *c*_2_, generate the primary undulatory waves underlying locomotion and are defined as:

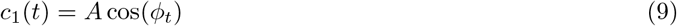

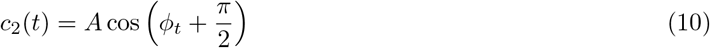

where the global amplitude is fixed at *A* = 7 and the second component is uniformly phase-shifted by *π/*2.

The temporal evolution of the phase variable is governed by a behavior-dependent angular frequency, *ω*(*y*_*t*_). The phase updates at each timestep according to:

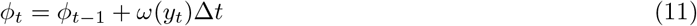

where Δ*t* = 1*/*32s corresponds to the temporal resolution of the reconstructed kinematic sequence (32 frames per second). This angular frequency dictates the speed and direction of the underlying undulatory wave. For standard forward locomotion, the angular frequency is set to *ω*_fwd_ = 1.14*π* rad/s. During slow forward locomotion, the frequency is attenuated to *ω*_slow_ = 0.6*π* rad/s. For all reverse states (aggregating “reverse 1”, “reverse 2”, and “reverse sus”), the wave direction is inverted by applying a negative angular frequency, *ω*_rev_ = −*π* rad/s.

When the model enters a turning state (“ventral turn” or “dorsal turn”), the first two coefficients continue to update using the forward locomotion phase step, while the third eigenworm coefficient, *c*_3_(*t*), is recruited to execute the turn. The raw turning signal is set to +4 for dorsal turns, −4 for ventral turns, and 0 otherwise. To prevent unrealistic, instantaneous postural twitches caused by discrete state transitions, this step-like *c*_3_(*t*) sequence is smoothed over time. Specifically, we convolve the *c*_3_(*t*) signal with a Gaussian temporal window (2 seconds in duration, truncated symmetrically at 3 standard deviations) to ensure smooth kinematic transitions into and out of turns.

Finally, having generated the continuous time series for the three coefficients, we reconstruct the relative angles between adjacent body segments by multiplying the coefficients by their respective eigenworm basis vectors. These relative angles are spatially resampled to define a uniform model comprising 25 discrete segments. By integrating these relative angles along the body axis, we calculate the cumulative, absolute spatial coordinates of the virtual worm segments, enabling the 2D visualization of the generated behavioral trajectories.

While the eigenworm projection yields a continuous sequence of postural configurations (a synthetic kymogram), these relative angles describe the worm’s shape in an ego-centric frame, effectively depicting the animal undulating “in place.” To translate these postural sequences into 2D spatial trajectories, we utilize the ElegansBot biomechanical simulation framework (Chung et al., 2024). The synthetic kymograms serve as the kinematic driving input for the ElegansBot model, which numerically simulates the dynamic physical interactions between the multi-segmented body and a virtual environment. The model transforms the relative postural changes into absolute, time-resolved spatial coordinates for each body segment within a global reference frame, yielding realistic 2D simulated trajectories of the virtual worm navigating its environment.

### 4.5 Latent manifold steering for goal-directed behavior

The base generative framework autonomously simulates spontaneous neural and behavioral activity. To transition from modeling spontaneous dynamics to enacting goal-directed control, we introduce a latent steering mechanism. This allows us to perturb the autonomous latent trajectory to reach specific behavioral targets without retraining the foundational generative manifold.

We formalize these goals by defining a set of desired, incentivized behaviors *Y*_*g*_ and a set of undesired, disincentivized behaviors *Y*_*b*_. Rather than imposing a complex prior over the entire trajectory, we introduce a deterministic steering module, *T*_*ψ*_, parameterized by a distinct set of neural network weights *ψ*. This steering network operates as a residual correction mechanism on the autonomous transition dynamics. At each generative timestep, the base transition model proposes a new state *z*_*t*_. The steering network takes both this proposed state and the preceding state *z*_*t*−1_ as inputs to compute a targeted intervention, yielding the steered latent state:

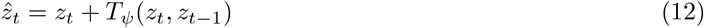

To optimize the steering parameters *ψ*, we freeze the weights of the base encoder, transition, and decoder models (*ϕ* and *θ*). We then perform long autoregressive rollouts starting from random initial conditions sampled from the learned latent manifold. The steering network is trained to shift the decoded behavioral probabilities toward *Y*_*g*_ and away from *Y*_*b*_ by minimizing the following objective:

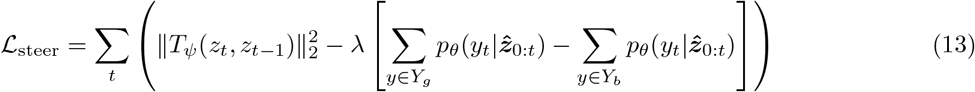

The objective function balances two competing forces. The terms scaled by *λ* drive the behavioral distribution toward the target sets using the differentiable logits of the behavioral decoder. Conversely, the *L*_2_ regularization term on the output of *T*_*ψ*_ penalizes the magnitude of the steering interventions. This regularization is critical: it ensures that the applied perturbations are as sparse and minimal as possible, forcing the steered trajectory to remain biologically plausible and bound to the natural dynamical manifold discovered during the initial training phase.

To ensure stable convergence and prevent the steering module from immediately forcing the trajectory off the manifold before finding valid dynamical paths, we employ a deterministic annealing schedule for the balancing parameter *λ*. Optimization begins with *λ* = 0, meaning the steering network initially learns to output zero vectors (acting as an identity mapping). The parameter is then linearly increased to a maximum value of *λ* = 2 over the first 50 training iterations, smoothly introducing the behavioral constraints into the optimization landscape.

## Acknowledgments

This research received funding from the European Research Council under the Grant Agreement No. 820213 (ThinkAhead), the Italian National Recovery and Resilience Plan (NRRP), M4C2, funded by the European Union – NextGenerationEU (Project IR0000011, CUP B51E22000150006, ‘EBRAINS-Italy’; Project PE0000013, ‘FAIR’; Project PE0000006, ‘MNESYS’), and the Ministry of University and Research, PRIN PNRR P20224FESY and PRIN 20229Z7M8N. The GEFORCE Quadro RTX6000 and Titan GPU cards used for this research were donated by the NVIDIA Corporation. We used a Generative AI model to correct typographical errors and edit language for clarity.

## A Supplementary Figures

**Figure S1:**
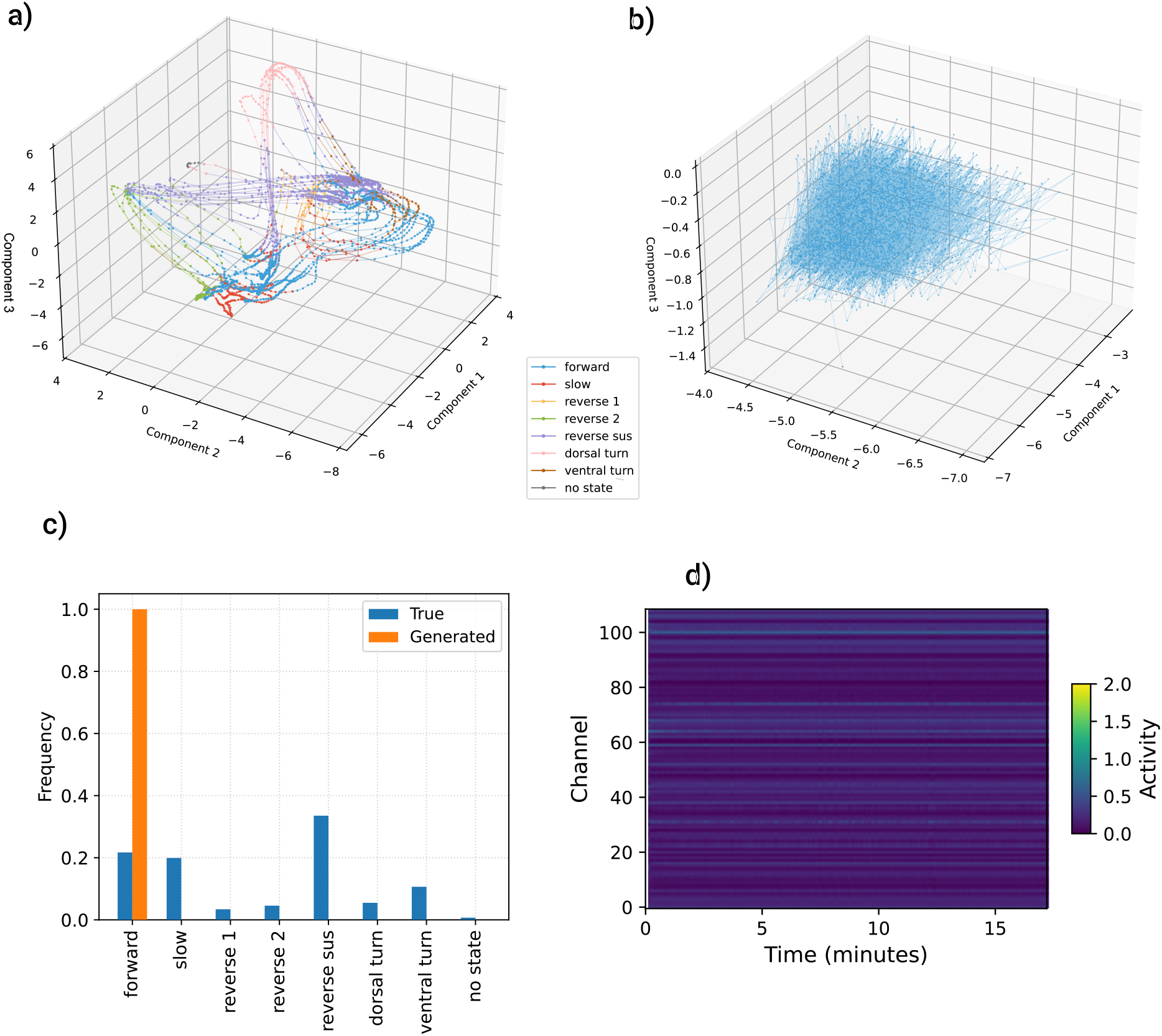
Ablation of recurrent temporal integration. We evaluated the necessity of temporal integration by restricting the training sequence window to a single transition step (containing only *x*_*t*_ and *x*_*t*+1_). **a)** Encoded manifold under temporal ablation. The training dataset represented within the three-dimensional latent space. A broad topological structure remains visible, but the distinct behavioral partitioning is noticeably degraded compared to the full recurrent model. **b)** Generative rollout failure. Latent trajectories produced by the transition module over 30,000 timesteps. Devoid of recurrent history to stabilize the dynamics, autonomous generation collapses into a localized, noisy distribution. **c)** Generative behavioral state frequencies. Comparison of behavioral frequencies demonstrating the complete failure of the generative process, with the collapsed latent trajectory decoded exclusively as “forward” locomotion with a probability of 1. **d)** Static generated neural activity. Heatmap showing the neural channels decoded from the generative rollout over 18 minutes. The collapsed, noisy latent dynamics result in completely static, featureless neural predictions across all channels.

